# Herpes Simplex Virus 1 Envelope Glycoprotein C Shields Glycoprotein D to Protect Virions from Entry-Blocking Antibodies

**DOI:** 10.1101/2024.08.20.608756

**Authors:** McKenna A. Hull, Suzanne M. Pritchard, Anthony V. Nicola

## Abstract

Herpes simplex virus 1 (HSV-1) gD interaction with the host cell receptor nectin-1 triggers the membrane fusion cascade during viral entry. Potent neutralizing antibodies to gD prevent receptor-binding or prevent gD interaction with gH/gL critical for fusion. HSV has many strategies to evade host immune responses. We investigated the ability of virion envelope gC to protect envelope gD from antibody neutralization. HSV-1 lacking gC was more sensitive to neutralization by anti-gD monoclonal antibodies than a wild type rescuant virus. gD in the HSV-1 gC-null viral envelope had enhanced reactivity to anti-gD antibodies compared to wild type. HSV-1 ΔgC binding to the nectin-1 receptor was more readily inhibited by a neutralizing anti-gD monoclonal antibody. HSV-1 ΔgC was also more sensitive to inhibition by soluble nectin-1 receptor. The viral membrane protein composition of HSV-1 ΔgC was equivalent to that of wild type, suggesting that the lack of gC is responsible for the increased reactivity of gD-specific antibodies and the consequent increased susceptibility to neutralization by those antibodies. Together, the results suggest that gC in the HSV-1 envelope shields both receptor-binding domains and gH/gL-interacting domains of gD from neutralizing antibodies, facilitating HSV cell entry.

**Importance:** HSV-1 causes lifelong infections. There is no vaccine and no cure. Understanding HSV immune evasion strategies is an important goal. HSV-1 gC is a multi-functional envelope glycoprotein. This study suggests that virion gC physically shields neighboring gD from antibodies, including neutralizing monoclonal antibodies. This mechanism may allow HSV to escape immune detection, promoting HSV infection in the host.

## Introduction

Herpes simplex virus 1 (HSV-1) is a ubiquitous pathogen that is estimated to affect 90% of adults worldwide (1). Typical symptoms include recurrent oral or genital lesions. Infection is lifelong and there is no vaccine (2). Grave outcomes of HSV infection include encephalitis, blindness, and disseminated infections of the immunocompromised (3, 4). The high prevalence and persistence of HSV is partly due to immune evasion strategies employed by the virus.

HSV-1 glycoprotein C (gC) is a multifunctional 511 amino acid, type 1 membrane glycoprotein present in the virion envelope and on the surface of infected cells (5). gC is specific to the alphaherpesviruses. Virion envelope gC functions in viral entry into host cells (6–9). gC also plays roles in immune evasion and has been a focus of HSV vaccine strategies (10–18). Virion gC protects gB from antibody-mediated neutralization (13, 15). Here we investigate the ability of gC to shield the HSV receptor-binding protein gD.

HSV-1 glycoprotein D (gD) is a 369 amino acid type I envelope glycoprotein (19). Host cell receptors nectin-1 and HVEM bind to the same face of gD, near the C-terminus of the gD ectodomain (Fig. 1), but at distinct sites (20–29). Binding of gD to a cognate receptor triggers the movement of the C-terminal extension, revealing receptor contact sites on the core. The receptor-triggered structural change in gD is thought to initiate the membrane fusion cascade by promoting interaction with gH/gL (30–36). HSV-1 gD is the major target of neutralizing antibodies and is a prime target for vaccine development (37, 38). MAbs to gD can block HSV entry by preventing binding to host cell receptors or can block fusion with no effect on receptor binding.

**Fig. 1.**
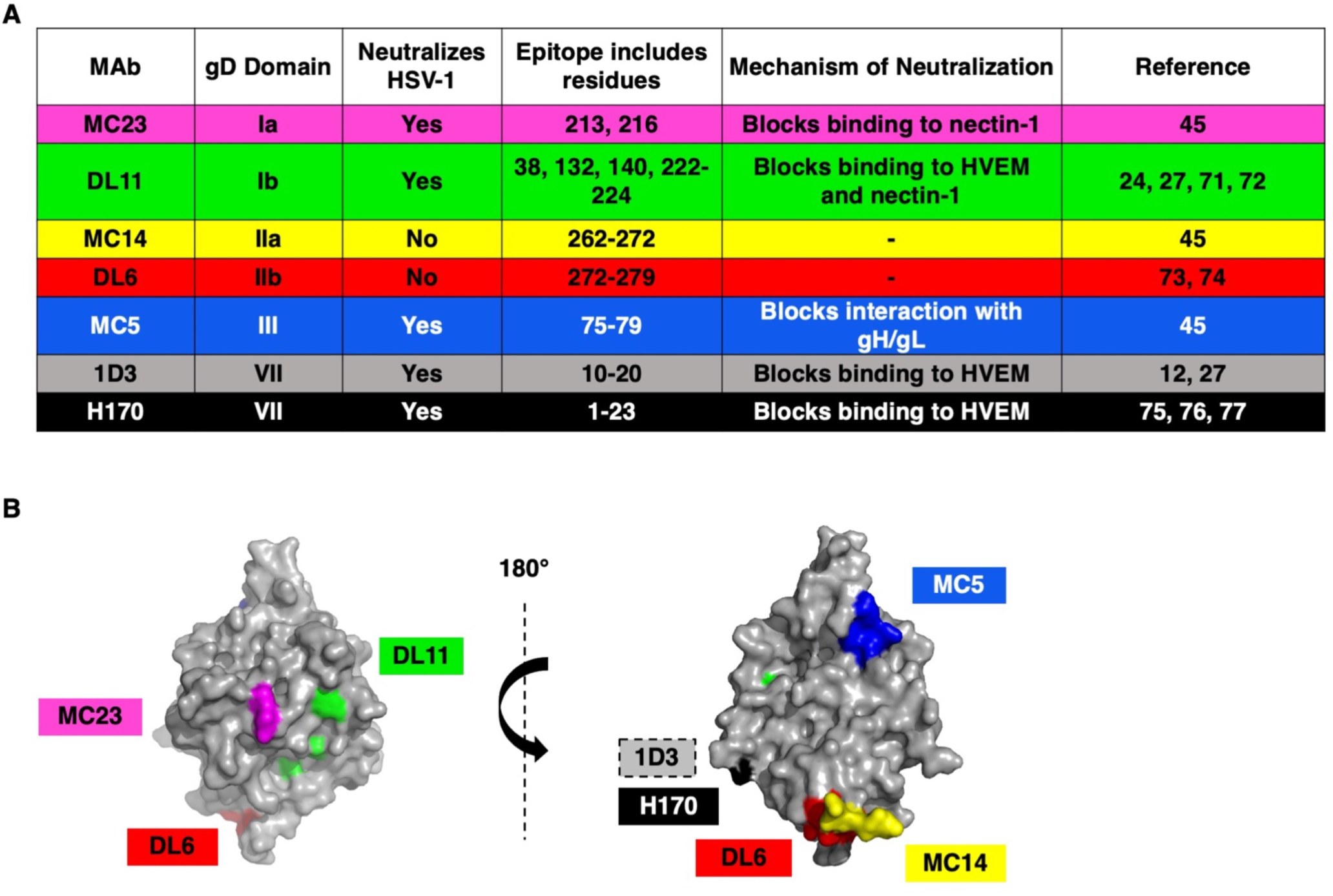
(A) Monoclonal antibodies to HSV-1 gD used in this study. (B) Structure of HSV-1 gD ectodomain (PDB accession number 2C36) (25) with MAb epitopes indicated. The receptor binding face of gD is on the left. MAb 1D3 binds to gD residues near the N-terminus that are not resolved in this structure.

In this study, we provide evidence that gC protects gD from antibody recognition of neutralizing epitopes. The envelope glycoproteins of several viruses protect themselves from antibody neutralization (39–42). The results support a unique viral immune protection mechanism whereby HSV-1 gC shields distinct neighboring glycoproteins from entry-blocking antibodies.

### The absence of gC renders HSV-1 more sensitive to neutralization by gD antibodies on two distinct cell types

To determine the impact of virion gC on HSV-1 infectivity in the presence of neutralizing MAbs we employed a panel of mouse anti-gD MAbs against multiple epitopes and functions of gD (Fig. 1). We tested two-fold dilutions of these MAbs ranging from 2 μg/mL to 9.76 x 10^-6^ μg/mL on Vero cells. HSV-1 neutralization was defined as a reduction in infectivity of >50% in the presence of anti-gD MAb. Importantly, HSV-1 gCR and ΔgC contain similar levels of viral proteins gB, gD, gH, and VP5 (data not shown) (9, 15). Thus, differences detected between the two viruses may be attributed to the lack of gC in the gC-null virus. HSV-1 ΔgC was more sensitive to MAb neutralization ranging from 2-16-fold more sensitive compared to HSV-1 gCR (Fig. 2). The negative control MAb MC14 failed to neutralize either virus, as expected (Fig. 2). MAb 1D3, which blocks gD from interacting with HVEM, neutralized ΔgC at a concentration of 3.9 x 10^-3^ μg/mL. 1D3 neutralized gCR at 0.125 μg/mL, which was 16-fold higher than the concentration required to neutralize ΔgC (Fig. 2E). MAb MC5, which blocks gD from interacting with gH/gL, neutralized HSV-1 ΔgC at 3.9 x 10^-3^ μg/mL and gCR at a concentration of 1.5 x 10^-2^ μg/mL on Vero cells (Fig. 2D). This was a 4-fold difference in MAb MC5 concentration. MAb MC23, which blocks gD interaction with nectin-1, required a 2-fold higher concentration to neutralize HSV-1 gCR. MAb DL11, which blocks gD interactions with both nectin-1 and HVEM, required an 8-fold higher concentration to neutralize gCR (Fig. 2A and B). In summary, 2-to 16-fold more antibody was required to neutralize HSV-1 when gC was present (Fig. 2G).

**Fig. 2.**
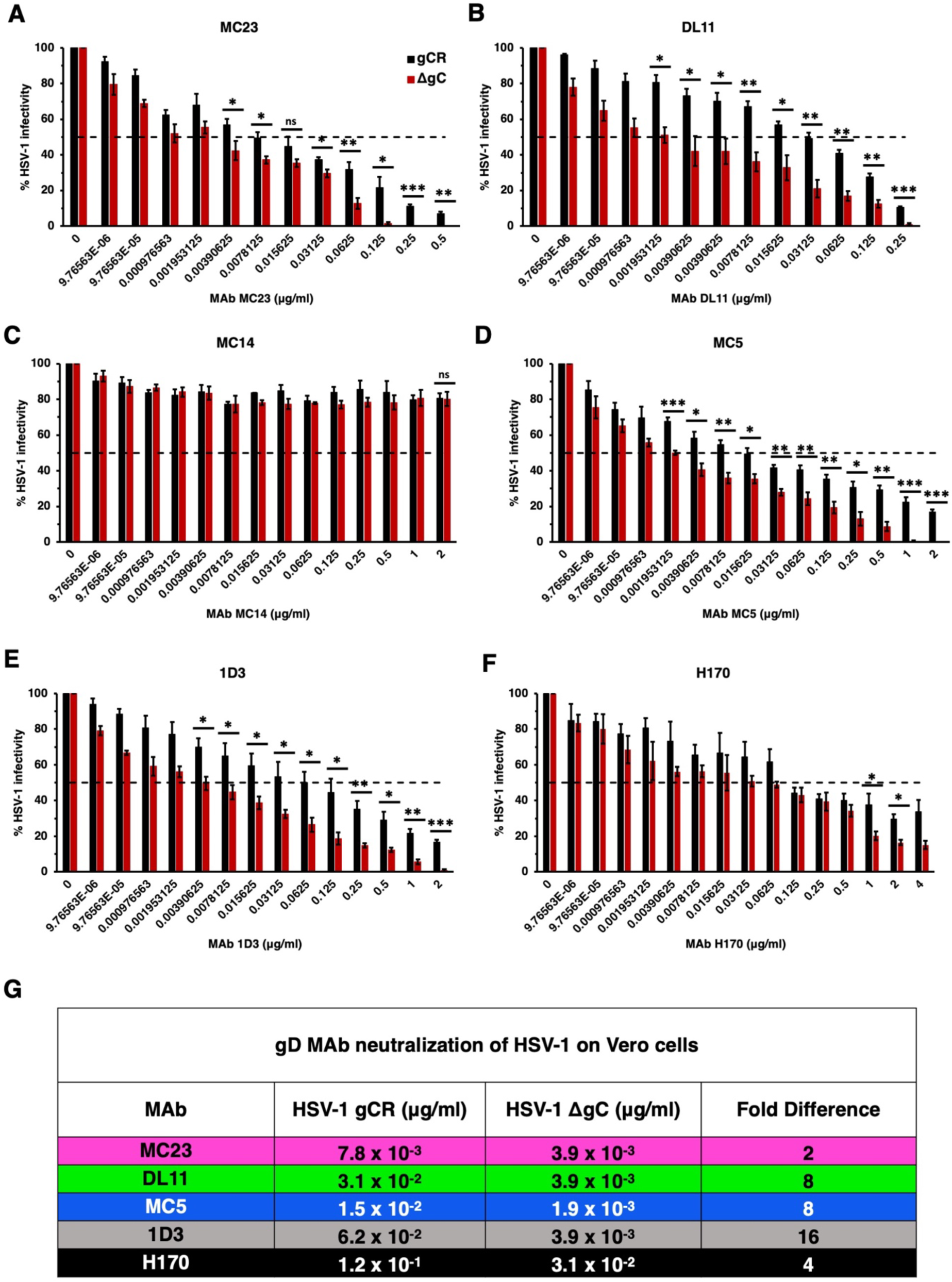
Neutralization of gC-null mutant HSV-1 infection of Vero cells by antibodies to gD. HSV-1 gCR (black) or HSV-1 ΔgC (red) (100 PFU) was treated with monoclonal antibodies MC23 (A), DL11 (B), MC14 (C), MC5 (D), 1D3 (E), or H170 (F) for 1 h at 37°C. Infectivity was determined by plaque formation on Vero cells. Each experiment was performed with triplicate samples. Values are the means and standard errors of results from three independent experiments. Statistical significance was determined via Student’s t-test where *, *p* < 0.05; **, *p* < 0.01; ***, *p* < 0.001; ns, not significant. (G) Antibody concentration at which > 50% of virus was neutralized. Fold difference was calculated by dividing the concentration of MAb required to neutralize HSV-1 gCR by the concentration of MAb required to neutralize ΔgC.

Next, we investigated whether virion gC impacts the ability of nectin-1 blocking antibodies to neutralize HSV-1 infection specifically mediated by nectin-1. Mouse melanoma B78 cells are resistant to HSV entry and must be supplied with a gD-receptor to render them permissive to HSV entry and infection (43). On B78-nectin-1 cells, MAb DL11 neutralized HSV-1 ΔgC at a concentration of 1.9 x10^-3^ μg/mL. Four-fold more DL11 (7.8 x 10^-3^ μg/mL) was required to neutralize gCR (Fig. 3B). MAb MC23 neutralized HSV-1 ΔgC at 7.8 x10^-3^ μg/ml and gCR at 0.3125 μg/ml. Thus, 4-fold more antibody was required to neutralize gCR (Fig. 3A). MAb 1D3 failed to neutralize HSV-1 infection mediated by nectin-1, as expected (Fig. 3C). In summary, on B78-nectin-1 cells, 4-fold more MAb was required to neutralize HSV-1 gCR compared to ΔgC (Fig. 3D). HSV-1 that lacks gC had enhanced sensitivity to anti-gD MAbs across two cell types and across all domains of gD tested. It was previously shown that MAb DL6 neutralized HSV-1 ΔgC at a dilution of 1:2000 on Vero cells and failed to neutralize gCR (15). These data demonstrate that the absence of gC renders HSV-1 more sensitive to neutralization by gD MAbs.

**Fig. 3.**
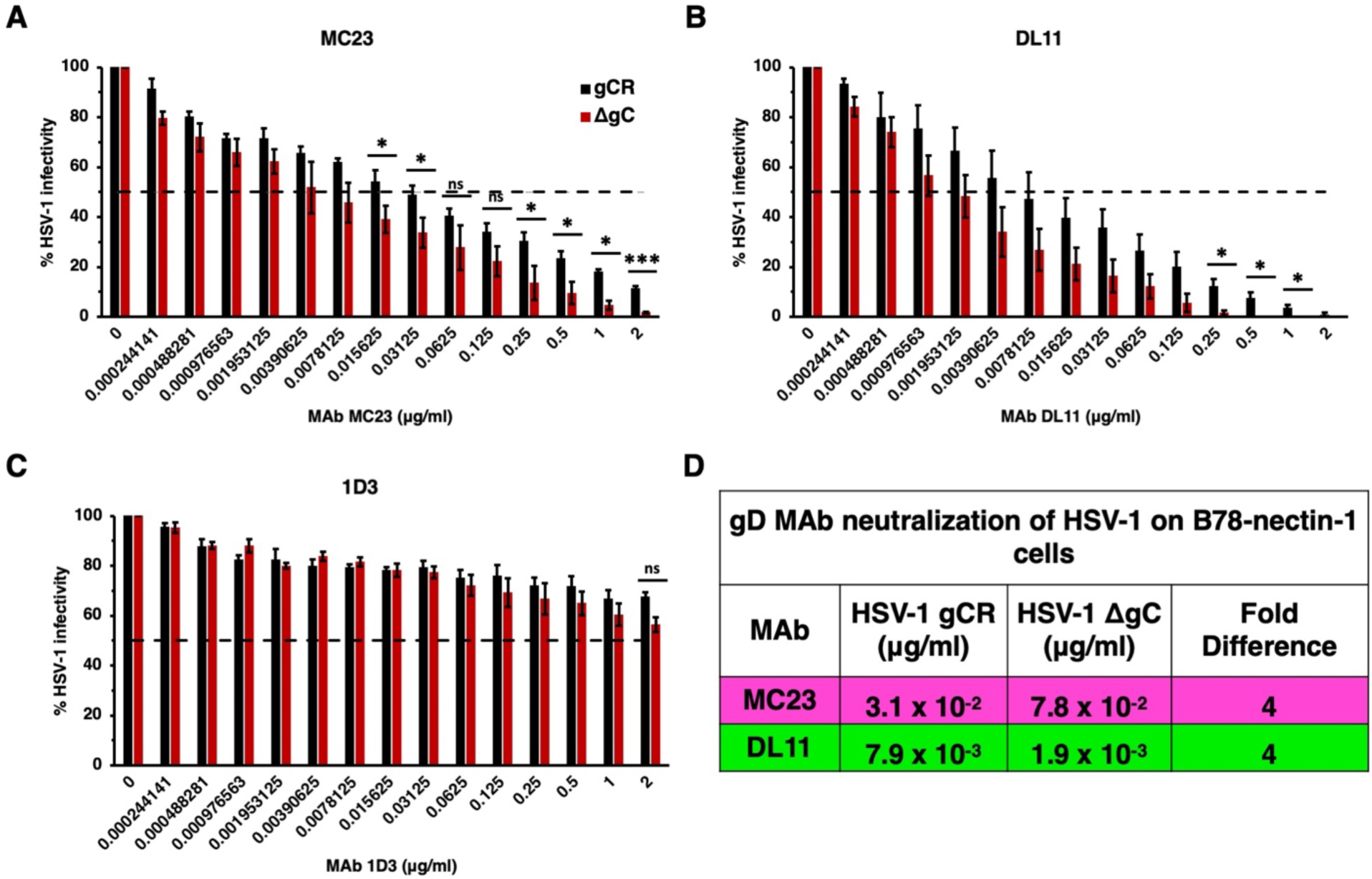
Neutralization of gC-null mutant HSV-1 infection mediated specifically by the nectin-1 receptor. HSV-1 gCR (black) or HSV-1 ΔgC (red) was treated with gD monoclonal antibodies MC23 (A), DL11 (B), or 1D3 (C) for 1 h at 37°C. Infectivity was determined by plaque formation on B78-nectin-1 cells. Values are the means and standard errors from three independent experiments. Statistical significance was determined via Student’s t-test where *, *p* < 0.05; **, *p* < 0.01; ***, *p* < 0.001; ns, not significant. (D) Antibody concentration at which > 50% of virus was neutralized. Fold difference was calculated by dividing the concentration of MAb required to neutralize HSV-1 gCR by the concentration of MAb required to neutralize ΔgC.

### The absence of virion gC enhances HSV-1 reactivity to gD antibodies

To interrogate the mechanism by which HSV-1 ΔgC is more sensitive to neutralization, we assessed the antigenic reactivity of gD MAbs with both HSV-1 gCR and ΔgC. The binding of the panel of gD antibodies to HSV-1 ΔgC and gCR was compared by dot blot immunoassay. Virus was blotted directly onto nitrocellulose membrane under native conditions. The membrane was probed with anti-gD Mabs, and binding was determined via fluorescence imaging (Fig. 4) followed by densitometry (Fig. 5). HSV-1 ΔgC was more sensitive to gD MAb binding compared to gCR, ranging from 2.7- to 5.6-fold more sensitive. MAb MC23, which blocks gD from interacting with nectin-1, bound to HSV-1 ΔgC 3.1-fold more intensely than to gCR (Fig. 4A and 5A). MAb MC14, which is non-neutralizing, bound to ΔgC 5.6-fold more intensely than to gCR (Fig. 4C and 5C). This trend remained constant across all domains of gD tested, with every antibody being more reactive with HSV-1 ΔgC. For ΔgC, there was an enhanced reactivity of 3.9-fold with DL11, 5.2-fold with DL6, 3.5-fold with MC5, 2.9-fold with 1D3, and 2.7-fold with 1D3 (Fig. 4 and 5). In summary, HSV-1 ΔgC was more sensitive to gD MAb binding regardless of the MAb’s epitope or function (Fig. 5H).

**Fig. 4.**
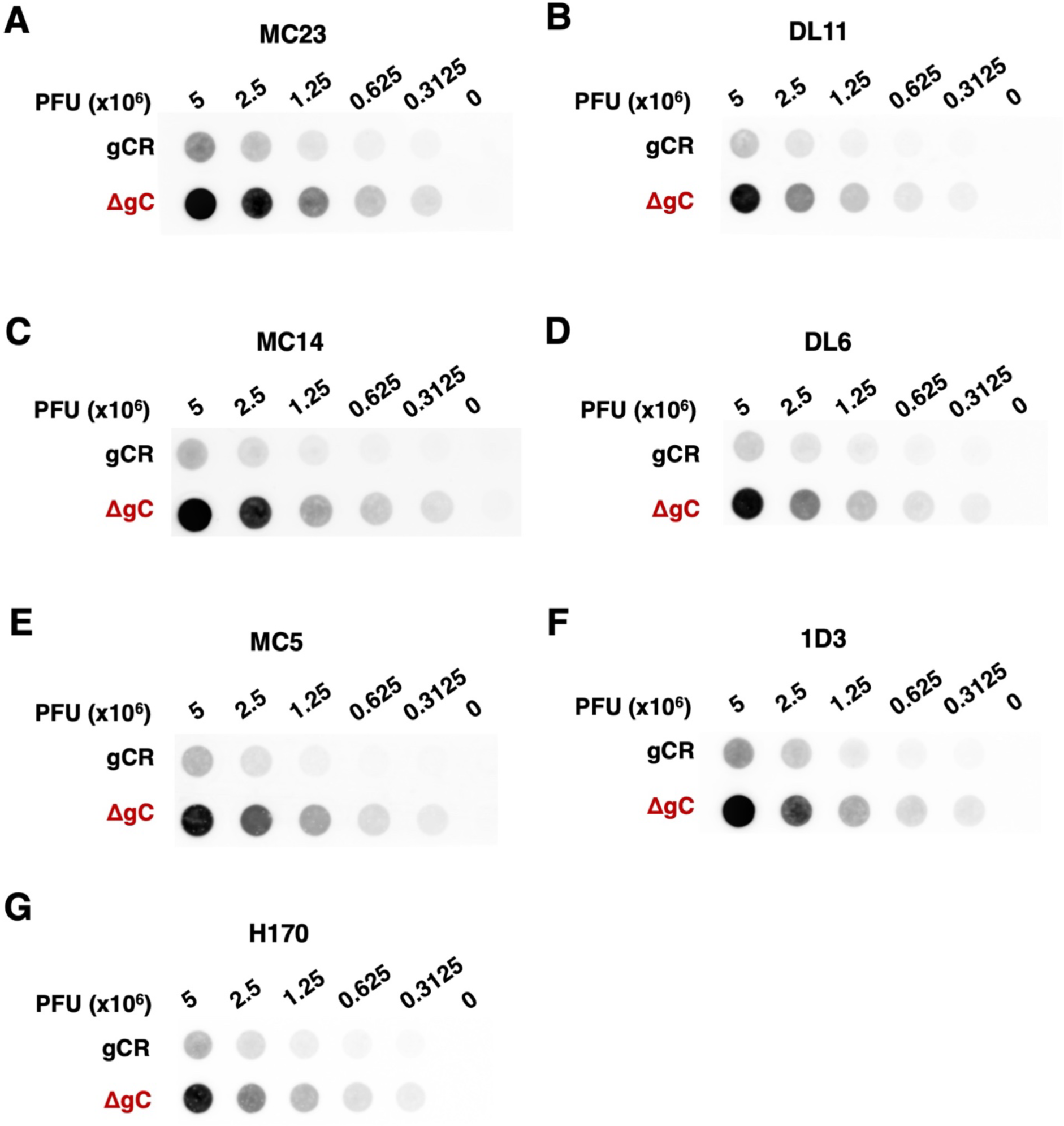
Reactivity of HSV-1 ΔgC with MAbs to gD. Equivalent infectious particles of HSV-1 ΔgC or gCR were serially diluted and blotted directly onto nitrocellulose membranes and probed with gD MAbs MC23 (A), DL11 (B), MC14 (C), DL6 (D), MC5 (E), 1D3 (F), or H170 (G). MAb reactivity was determined via densitometry with ImageJ (Fig. 5).

**Fig. 5.**
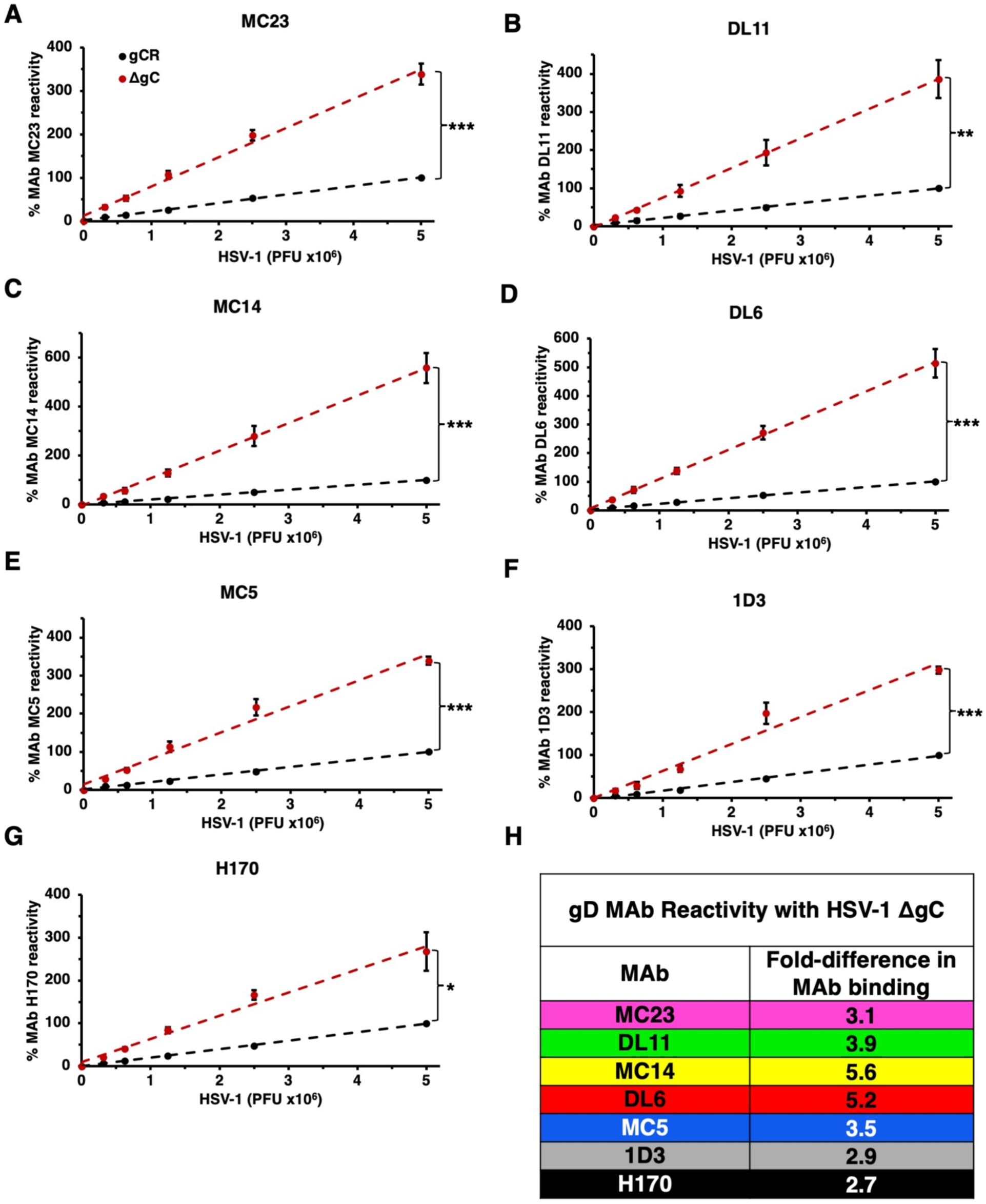
Reactivity of HSV-1 ΔgC with MAbs to gD. (A-G) HSV-1 ΔgC (red) or gCR (black) was blotted onto a nitrocellulose membrane and probed with antibodies against gD. Antibody reactivity was determined via densitometry with ImageJ. Results are the mean and standard error of three independent experiments. Representative blots are shown in Figure 4. Statistical significance was determined via Student’s t-test where *, *p* < 0.05; **, *p* < 0.01; ***, *p* < 0.001. (H) Differences in HSV-1 ΔgC and gCR reactivity were determined by comparing slopes of the best fit lines in panels A-G.

### The presence of gC in HSV-1 reduces the ability of a gD monoclonal antibody to block receptor binding

We next assessed the effect of gC on the ability of anti-gD antibody to inhibit HSV-1 binding to receptor. Soluble ectodomain forms of gD-receptors bind directly to HSV particles and block entry and infection (24, 27, 44). We tested the ability of HSV-1 ΔgC to bind to soluble nectin-1 in the presence of nectin-1 blocking gD MAb DL11 (24, 27). HSV-1 ΔgC or gCR was pre-incubated with gD MAb, and then soluble nectin-1 was added. Samples were layered onto a sucrose gradient and separated by ultracentrifugation. The virion fraction was recovered and blotted directly onto a nitrocellulose membrane, and then probed for the presence of soluble nectin-1 (Fig. 6A).

**Fig. 6.**
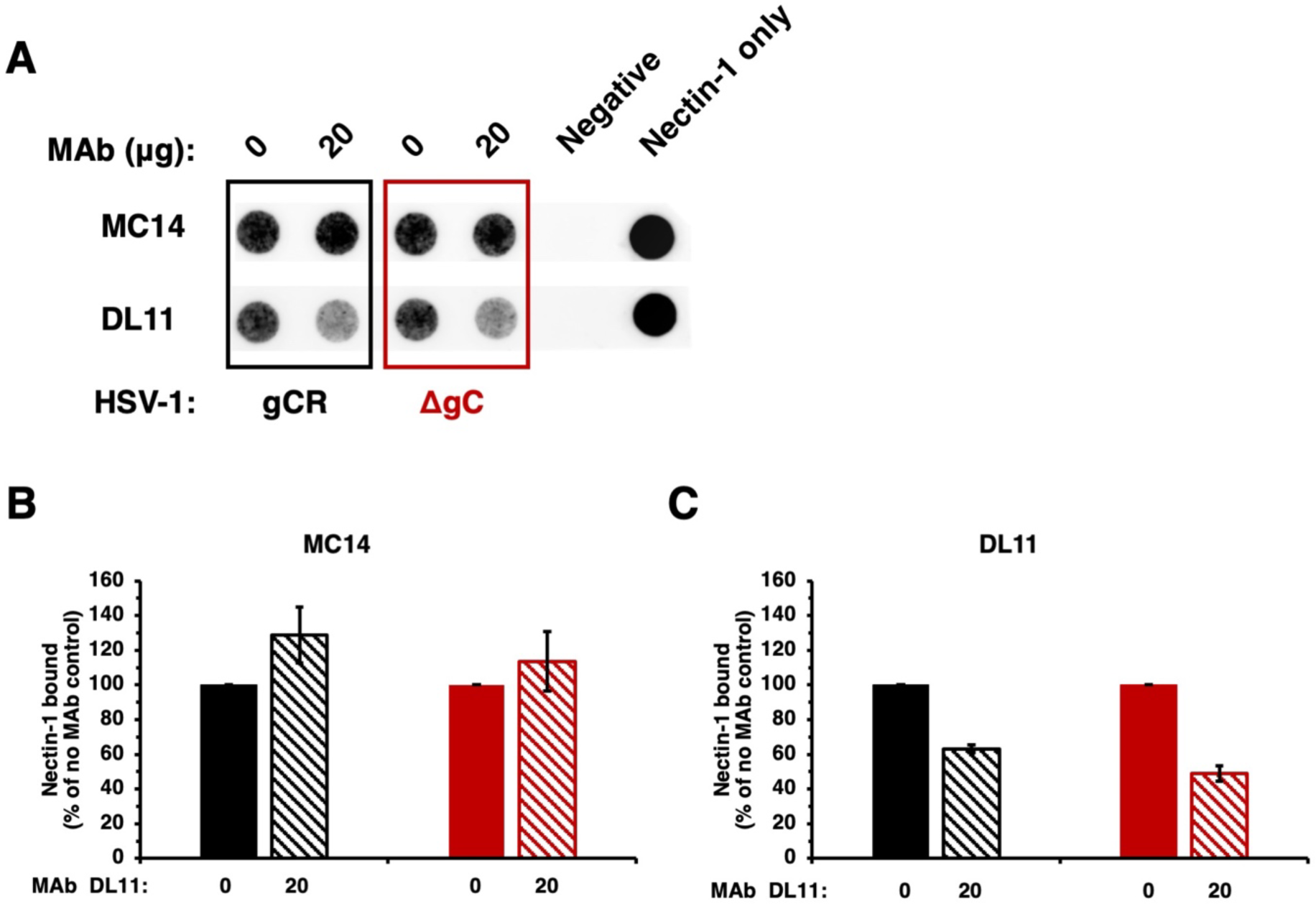
Inhibition of HSV-1 ΔgC binding to nectin-1 by gD antibodies. (A) HSV-1 ΔgC (red) or gCR (black) was treated with gD MAb DL11 or MC14 at 37°C for 1 h. Soluble nectin-1 was added at 4°C for 2 h. Samples were separated on a sucrose gradient, and the HSV-1-containing fraction was blotted onto a nitrocellulose membranes and probed with anti-6x-HIS tag MAb to detect nectin-1. (B, C) Nectin-1 binding was determined via densitometry with ImageJ. Results are the mean and standard error of three independent experiments.

Following pre-incubation with 20 μg MC14, a non-neutralizing gD MAb, soluble nectin-1 binding to HSV-1 ΔgC and gCR was not inhibited, as expected (Fig. 6A and B). MAb MC14 enhanced nectin-1 reactivity with both HSV-1 gCR and HSV-1 ΔgC, as previously reported (45). MAb MC14’s impact on ΔgC binding to nectin-1 was similar to gCR (Fig. 6B). gD MAb DL11 inhibited nectin-1 binding to both viruses (Fig. 6A). MAb DL11 inhibited 51% of soluble nectin-1 binding to HSV-1 ΔgC and inhibited 37% of soluble nectin-1 binding to gCR (Fig. 6C). This is consistent with findings from the dot blot assay (Fig. 4 and 5) and neutralization assay (Fig. 2 and 3).

### The absence of gC renders HSV-1 more sensitive to inhibition by soluble nectin-1

We investigated the ability of a recombinant nectin-1 ectodomain to block entry of HSV-1 in the absence of gC. Soluble nectin-1 receptor inhibits HSV-1 entry and infection by competing with receptors on target cells (44). To evaluate the ability of soluble nectin-1 to inhibit HSV-1 ΔgC entry, we conducted a β-galactosidase reporter assay. B78-nectin-1 cells contain the *E. coli lacZ* gene under the control of the HSV-1 ICP4 gene promoter (43). HSV-1 ΔgC or gCR was incubated with soluble nectin-1 for 2 h at 4°C and then added to B78-nectin-1 cells. At 6 h p.i., β-galactosidase activity was determined. Soluble nectin-1 inhibited entry of both HSV-1 ΔgC and gCR in a concentration-dependent manner starting at 0.01 μM (Fig. 7). However, soluble nectin-1 hampered HSV-1 ΔgC entry more robustly than HSV-1 gCR. Following pretreatment with 1 μM soluble nectin-1, HSV-1 ΔgC entry was reduced to 21% vs. 48% for HSV-1 gCR entry (Fig. 7). Together, the results suggest that virion gC renders HSV-1 less sensitive to inhibition by both gD antibodies and soluble receptor.

**Fig. 7.**
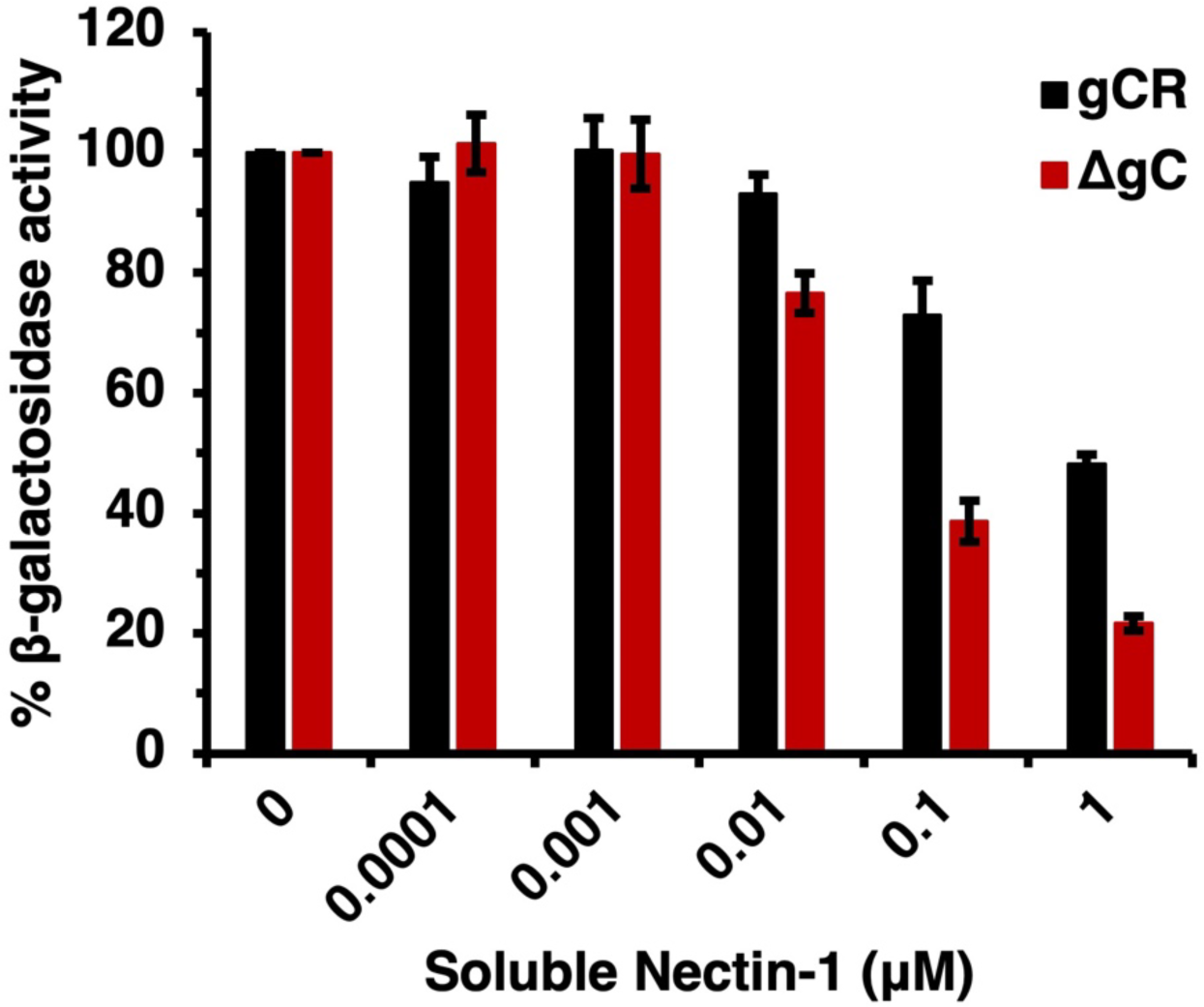
Inhibition of HSV-1 ΔgC entry by soluble nectin-1. HSV-1 ΔgC (red) or gCR (black) (2 x 10^5^ PFU) was treated with soluble nectin-1 at 4°C for 2 h and then added to B78-nectin-1 cells (MOI 5). At 6 h p.i., β-galactosidase activity was detected as an indicator of entry and infection. Results are the mean and standard error of three independent experiments.

## Discussion

HSV-1 gC has multiple functions during the viral infectious cycle, including entry, egress, and immune evasion. The current study demonstrates the ability of virion gC to shield the essential receptor-binding protein gD. We present several lines of evidence suggesting that gC in the HSV particle protects against neutralizing antibody to gD and against inhibition by soluble gD-receptors. We propose that gC is broadly shielding the entire neighboring gD molecule, including important functional domains for fusion and entry.

HSV-1 harbors many immune protective features that contribute to persistence in the host. HSV-1 gE, an envelope glycoprotein that is non-essential for entry, forms a high affinity Fc receptor with its partner gI. gE/gI binds to the Fc region of immunoglobulin G (IgG) antibodies to prevent epitope recognition (46, 47). gC prevents complement activation by binding and sequestering complement protein C3b (12, 18). Antibodies to gC can block this function (48). The increased sensitivity of gC-null HSV-1 to antibody-mediated neutralization (Fig. 2 and 3) can be explained at least in part by enhanced binding of antibodies to the virus in the absence of virion gC (Fig. 4 and 5). Neutralizing and non-neutralizing antibodies bound better to HSV that lacks gC. gC also shields gB and gH/gL from monoclonal antibody binding and neutralization (13, 15). This protective role is specific to gC. The absence of gE from the HSV-1 particle had little to no effect on MAb-mediated neutralization of HSV-1 (15).

For several viruses including influenza, HIV, and Nipah virus, the N-linked glycans of the viral fusion protein shield its own epitopes from neutralization (39–42, 49). The N-linked glycans of HSV-1 fusion protein gB provide self-protection against antibody-mediated neutralization and antibody-dependent cytotoxicity (50). gC is not a fusion protein, but it contains a heavily glycosylated N-terminal domain. Future research will determine whether N-glycans on gC shield neighboring glycoproteins. Whether N-glycans on gC block the binding of anti-gC antibodies also remains to be determined. This would be a unique feature for a non-fusion glycoprotein. Importantly, gC is in close enough proximity to gD to be chemically crosslinked in HSV particles (51). However, direct interaction between gC and gD has not been detected. Low-affinity or transient interactions may be difficult to capture. Physical interactions between and among HSV-1 gD, gH/gL and gB have also been difficult to capture, despite demonstrations of functional interactions (33, 52–56). The specifics of how gC protects neighboring glycoproteins from antibody-mediated neutralization is the subject of ongoing work.

Initial attachment of HSV-1 to the host cell is mediated by gC interaction with cell surface proteoglycans, principally heparan sulfate (6, 7). Alphaherpesviruses utilize low pH endosomal entry pathways in a cell-specific manner (57–62). The fusion protein gB undergoes well-documented antigenic changes upon exposure to mildly acidic pH, such as that present in the host cell endosomes (63–68). During endosomal entry into epithelial cells, gC undergoes pH-triggered changes and is thought to regulate the conformational change and function of the fusion protein gB (8, 9). gC also enhances virion release from infected cells (69).

This study highlights gC as an immune protective molecule that shields neighboring entry glycoproteins from neutralizing antibody binding and activity. Antibodies against gC can block immune evasion functions (14). Several vaccine candidates for HSV-1 and HSV-2 contain two or more different surface glycoprotein immunogens, including gC (10, 11, 14, 16, 17). Inclusion of gC in an HSV vaccine may block intrinsic protective properties of HSV.

## Materials and Methods

### Cells and viruses

Vero cells (American Type Culture collection; ATCC; Rockville, MD) were cultured in Dulbecco’s modified Eagle’s medium (DMEM; Life Technologies Corporation, Grand Island, NY) supplemented with 10% fetal bovine serum (FBS; Atlanta Biologicals, Atlanta, GA) and penicillin, streptomycin, and glutamine (PSG; Life Technologies Corporation). B78 murine melanoma cells expressing nectin-1 (B78-nectin-1) (43), gifted by G. Cohen and R. Eisenberg (University of Pennsylvania), were cultured with the same medium. B78-nectin-1 cells were selected every third passage in culture medium supplemented with 250 μg/mL geneticin (G418; Sigma-Aldrich, St. Louis, MO) and 6 μg/ml puromycin (Sigma-Aldrich). HSV-1 KOS strain with the gC gene deleted, HSV-1ΔgC2-3 (ΔgC), and its rescuant, HSV-1gC2-3R (gCR) (70) were gifts from C.R. Brandt (University of Wisconsin, Madison).

### Antibodies

Anti-HSV-1 gD mouse monoclonal antibodies MC23 (domain Ia) (45), DL11 (domain Ib) (24, 27, 71, 72), MC14 (domain IIa) (45), DL6 (domain IIb) (73, 74), MC5 (domain III) (45), and 1D3 (domain VII) (12, 27) were gifts from G. Cohen and R. Eisenberg (University of Pennsylvania). H170 (domain VII) (75–77) was purchased from Virusys (Milford, MA).

### Plaque inhibition (neutralization) assay

Antibodies to gD were diluted two-fold in complete DMEM to achieve final concentrations ranging from 2 μg/ml to 2.4 x 10^-4^ μg/ml. HSV-1 ΔgC or gCR (100 PFU) was added to the antibody dilutions and incubated at 37°C for 1 h. The antibody-virus mixture was added to subconfluent Vero cells or B78-nectin-1 cells grown in 24-well plates. At 1 h p.i., the antibody-virus mixture was removed and replaced with fresh culture medium. At 18 to 24 h p.i., cells were fixed with an ice-cold 1:2 methanol-acetone solution. Plaque formation was determined by immunoperoxidase staining. Anti-HSV polyclonal antibody HR50 (Fitzgerald Industries International Inc., Acton, MA) was added to cells overnight at room temperature. Pro A-horseradish peroxidase (Invitrogen, Rockford, IL) secondary antibody was added for 2 h at room temperature. 4-chloro-1-napthol substrate (Sigma-Aldrich) was added for 15 min at room temperature. A MAb was considered neutralizing if there was a >50% reduction in plaque formation (infectivity).

### Dot blot assay

Serial dilutions of cell-free HSV-1 ΔgC or gCR, were prepared in Dulbecco’s phosphate buffered saline (PBS)(Life Technologies Limited, Paisley, UK). Samples were blotted onto a nitrocellulose membrane using a Minifold dot blot system (78) (Whatman, Kent, UK). Five percent milk in 0.2% PBS-Tween 20 blocking buffer was added, and the membrane was gently rocked for 30 min. Primary anti-HSV-1 gD antibody was prepared in blocking buffer and added to the membrane overnight at 4°C. Goat-anti-mouse polyclonal antibody conjugated with Alexa Fluor 647 (Invitrogen) was prepared in blocking buffer and added to the membrane at room temperature for 30 min. The membrane was imaged with an Azure Biosystems c400 fluorescent western blot imager and quantified via densitometry (ImageJ).

### Receptor binding assay

VP5 equivalents of HSV-1 ΔgC or gCR were incubated with 20 μg anti-gD-MAbs MC14 or DL11 in 10% BSA in PBS for 1 h at 37°C. 15 μg of a soluble ectodomain form of nectin-1 (containing amino acids Gln 31 – Thr 334) truncated prior to the transmembrane region and containing a C-terminal 6 x His tag (ACRO Biosystems, Newark, DE) was added. The mixture was incubated at 4°C for 2 h. Samples were added to the top of a 60%-30%-10% sucrose/PBS gradient and centrifuged at 16,000 x *g* for 4.5 h at 4°C with an SW32 Ti rotor (Beckman, Brea, CA). Virus bands at the 60%-30% sucrose interface were collected via tube side puncture. Virus bands were then blotted onto nitrocellulose membrane. Membranes were incubated in blocking buffer as described above. To detect nectin-1, a 6x-HIS antibody conjugated with CoraLite Plus 647 (Proteintech Group, Rosemont, IL) was added for 1.5 h at RT. The membrane was imaged with an Azure Biosystems c400 fluorescent western blot imager and quantified via densitometry (ImageJ)

### β-galactosidase reporter assay for HSV-1 entry

HSV-1 gCR or ΔgC was incubated with 1 x 10^-4^ μM to 1 μM soluble nectin-1 in cell culture medium for 2 h at 4°C. B78-nectin-1 cells were infected with the virus-nectin-1 mixture in quadruplicate for 6 h at 37°C. Cells were lysed with 1% IGEPAL C-630 (Sigma-Aldrich) and frozen at -80°C overnight. Lysates were thawed and chlorophenol red-beta-d-galactopyranoside (Roche Diagnostics, Indianapolis, IN) substrate was added. β-galactosidase activity was read at 595 nm with an ELx808 microtiter plate reader (BioTek Instruments, Winooski, VT).

## Acknowledgments

We thank Curtis Brandt, Gary Cohen, and Roselyn Eisenberg for the gifts of reagents. This work was supported by National Institutes of Health grant R56AI119159 (A.V. N.).

